# Strain-Specific Surface Polysaccharides Mediate Bacterial Induction of Metamorphosis in the coral *Pocillopora acuta*

**DOI:** 10.1101/2025.08.22.671858

**Authors:** Marnie L Freckelton, Brian T Nedved, Michael G Hadfield

## Abstract

Coral-reef ecosystem persistence depends on successful larval recruitment, a process increasingly jeopardized by climate change and other anthropogenic stressors. Microbial biofilms induce larval settlement and metamorphosis in marine invertebrates, including corals, but specific molecular cues for corals remain unclear. Glycosylated lipids and polysaccharides from crustose coralline algae (CCA), along with bacterial small molecules, are proposed inducers, but their effects vary by coral species, bacterial strain, and environment, highlighting the need to clarify their molecular basis and specificity. Here, we identify bacterial lipopolysaccharide (LPS), specifically its O-antigen polysaccharide component, as a potent settlement cue for larvae of the coral *Pocillopora acuta*. Using purified LPS from diverse marine bacteria, we show that inductive activity is bacterial species and strain specific and correlates with the structure of O-antigen. These findings support a glycan-centred mechanism of coral recruitment, implicating microbial surface chemistry as a key determinant of larval settlement. By demonstrating a specific and functional role for O-antigen, we expand the ecological relevance of bacterial glycans beyond immunity and pathogenesis, highlighting their role in key developmental and ecological transitions in marine invertebrates. As coral reefs face escalating environmental pressures, employing such signals offers a promising strategy to enhance larval settlement and support reef restoration.

## Introduction

Coral reefs are among the most biodiverse and ecologically critical ecosystems on the planet, supporting approximately 25% of all marine species and providing coastal protection, fisheries, and income for over half a billion people (1,2). Yet these ecosystems face intensifying threats from climate change, habitat degradation, overfishing, and pollution (2). Without rapid, coordinated, and sustained intervention, projections suggest that most coral reefs could be lost by 2050 (3,4). As restoration initiatives gain urgency, their chances of long-term success can be improved by understanding and leveraging the critical early life stage at which coral larvae are recruited to the reef (5–7).

Most marine invertebrates, including corals, begin life as planktonic larvae that must undergo settlement and metamorphosis to become benthic adults (8–10). These processes are rapid and irreversible, often unfolding within minutes to hours once specific environmental and biochemical conditions are encountered; they result in a permanent transition to a benthic lifestyle (8–10). The outcome of this event determines not only the fate of individual larvae but also patterns of population genetic structure and species distribution (8). As such, identifying the molecular nature of these cues is essential for understanding larval recruitment and for designing tools to enhance settlement in reef restoration efforts.

Corals as ecologically important species have been the subject of significant focus into the identification of cues for settlement and metamorphosis. Much of this research has focused on acroporid corals and their interactions with crustose coralline algae (CCA), particularly *Porolithon onkodes*, which is widely regarded as a potent inducer of settlement (11,12). Numerous studies have shown that larvae of *Acropora* spp. reliably respond to live CCA and its biochemical extracts, including ethanol- and water-soluble fractions, suggesting that specific algal morphogens play a key role in initiating metamorphosis (12–16). Tebben et al. (13) identified glycoglycerolipids and unresolved polysaccharides in their CCA fractions that induced settlement and metamorphosis in the larvae of *Acropora tenuis*. However, findings from Whitman et al. (12) challenge the assumption that CCA universally induces settlement across coral taxa. They demonstrate that while live fragments of *P. onkodes* induce strong responses in *Acropora spp*., larvae of many non-acroporid corals either responded weakly or not at all.

Alongside the focus on CCA, a growing body of research has explored the role of bacterial biofilms in inducing coral settlement. Indeed, a number of corals have been found to respond to bacterial biofilms without the presence of CCA, including *Acropora microphthalma* (17), *Acropora millepora* (18), *Acropora palmata, Orbicella franksi, Porites asteroides* (19), *Pocillopora (damicornis) acuta* (20), *and Leptastrea purpurea* (21). To date, two cues from bacteria have been isolated and identified, tetrabromopyrrolle (TBP) (19,22) and cycloprodigiosin (23), both isolated from *Pseudoalteromonas* species. While TBP has been shown to induce settlement in multiple coral species, including *Acropora palmata, Orbicella franksi*, and *Porites astreoides* (19), its ecological relevance remains debated (13). Its natural abundance on reefs is low, and while it can induce metamorphosis, it frequently fails to promote larval attachment, suggesting that it may function as a partial or context-dependent cue (13).

Outside of coral taxa, bacterial biofilm induced metamorphosis has been demonstrated in representative taxa from every major marine invertebrate phylum with pelagic larvae—including molluscs, annelids, echinoderms, crustaceans, and sponges (10,24). In the course of exploring potential settlement sites, coral larvae, as all marine larvae, encounter diverse microbial biofilms (10,24). These bacterial biofilms form rapidly on all submerged surface, abiotic and biotic, and their composition varies with substrate type and local environmental conditions (25). This variation generates spatially and temporally dynamic chemical landscapes with the potential to signal the suitability of an environment for continued growth and development (26–28). Insights into the molecular mechanisms behind bacterial induction of settlement and metamorphosis in corals, may benefit from other invertebrate model systems such as the polychaete tubeworm, *Hydroides elegans*. A cosmopolitan member of the biofouling community, *H. elegans* can be induced to settle and metamorphose by some, but not all, biofilm bacteria (29–31) and will not settle without the presence of bacteria (32,33). Recently, it was discovered that lipopolysaccharide (LPS), specifically its polysaccharide (O-antigen) component, is responsible for inducing metamorphosis in larvae of *H. elegans* in response to the bacterium *Cellulophaga lytica*. LPS, which is produced by Gram-negative bacteria and located in the outer leaflet of the outer membrane, exhibits structural variability that is influenced by both bacterial taxonomy and environmental conditions, making it a promising candidate for a broadly relevant yet environmentally tunable metamorphic cue. This result is in keeping with an emerging body of research that supports the hypothesis that surface-associated glycans represent a common biochemical motif among settlement-inducing cues for marine invertebrate larvae from both algal and bacterial sources (13,34–36).

Notably, some of the same bacterial species shown to induce tubeworm metamorphosis are also found on coral reef substrates and have been reported to induce settlement in *P. acuta* (20). However, the inductive potential of their LPS has not been tested directly, and it remained unknown whether LPS alone can explain biofilm-mediated settlement in *P. acuta*. In this study, we evaluated whether LPS from coral reef-associated bacterial strains can induce metamorphosis in larvae of *P. acuta* in the absence of live biofilms or algal cues. We focused on biofilm bacterial species previously isolated from Hawaiian coral reefs and identified by Tran and Hadfield (20) to be inductive for larvae of *P. acuta*. Specifically, we asked: (i) is purified LPS sufficient to trigger settlement; (ii) does the inductive capacity vary among LPSs from different bacterial strains; and (iii) do structural features of the O-antigen polysaccharide mediate the larval response.

To address the third question, we used lectin-binding assays to characterize terminal sugar residues on each LPS. Lectins, widely used in other invertebrate systems, provide insight into glycan structure and potential larval recognition motifs (37,38). To further test whether inductive activity depends on the polymeric structure of the O-antigen, we also exposed larvae to individual monosaccharides corresponding to lectin-binding residues. This allowed us to assess whether specific sugar identities alone are sufficient, or whether higher-order glycan structure is required for inducing settlement.

By linking known settlement cues across phyla and testing the role of LPS in *P. acuta*, this study advances our understanding of coral–microbe interactions and contributes to the broader recognition of surface-bound polysaccharides as conserved mediators of larval settlement. We hypothesized that lipopolysaccharide (LPS), particularly its structurally diverse O-antigen component, represents a conserved microbial cue capable of inducing settlement and metamorphosis in larvae from across a broad range of marine invertebrate phyla. The experimental evidence presented in this study supports this hypothesis, indicating that such a mechanism may be shared across taxa and suggesting the existence of a fundamental, evolutionarily conserved pathway underlying larval recruitment in the seas (24). Building on this understanding, microbial tools leveraging such cues may be designed to enhance coral settlement, offering a biologically grounded approach to support reef restoration efforts (39).

## Methods

### Collection and culture of larvae from *Pocillopora acuta*

Adult colonies (10 cm) of the brooding coral *Pocillopora acuta* were collected from Kane"ohe Bay, Hawai"i. *P. acuta* released larvae on the full moon and larvae were competent to settle within 7 days. Larvae were cultured to competency in filtered seawater at a concentration of 1 larva/ml, at 27°C with daily water changes and a 12 h light-dark cycle. Historically both *P. damicornis* and *P. acuta*, two species that have been difficult to distinguish morphologically (40) have been identified within Kāne"ohe Bay, Hawai"i. To ensure that the collected corals were solely *P. acuta*, we sequenced the hypervariable mitochondrial ORF to genotype all colonies according to the method of Flot et al. (41).

### Bacterial culture

Gram-negative bacteria isolated from biofilms in Hawai"i and employed in these studies were *Cellulophaga lytica* HI1 isolated from within the port of Pearl Harbor, Hawai"i (31,42), *Pseudoalteromonas luteoviolacea* H1 isolated from within the port of Pearl Harbor, Hawai"i (31,43), *Pseudoalteromonas luteoviolacea* ATCC 33492 isolated from surface seawater in France (44,45), *Pseudoalteromonas luteoviolacea* B1P isolated from Kane"ohe Bay, Hawai"i, Hawai"i (20), *Thalassotalea euphylliae* MR31e isolated from Kane"ohe Bay, Hawai"i, Hawai"i (20), *Thalassotalea euphylliae* M23b isolated from Kane"ohe Bay, Hawai"i, Hawai"i (20), and *Tenacibaculum aiptasiae* T48 isolated from within the port of Pearl Harbor, Hawai"i (34). Bacteria were streaked from −80°C glycerol stocks onto half seawater tryptone agar (1/2FSWt) (46) and incubated at 25°C for 24-48 hours. Single colonies were used to inoculate 3ml broth cultures and incubated for 4 hours at 28°C with shaking (170 rpm). These starter cultures were adjusted to an OD_600_ of 1.000, and aliquots were used to inoculate overnight cultures. Cells were harvested by centrifugation (4,000 *g*, 30 min, 4°C) and washed with 1/10th volume of double-filtered autoclaved seawater (DFASW). Bacterial cells were then either frozen for later LPS extraction or diluted for biofilm formation.

### Monospecific biofilms

Washed bacterial cells were resuspended in DFASW and adjusted to produce a cell density of 10^8^ cells/ml for all strains (47). Adjusted bacterial solutions were added to 24-well plates and incubated to enable the attachment of cells. After 1 h, biofilmed wells were gently washed three times with DFASW to remove unattached cells. Biofilms were then ready for settlement assays.

### LPS extraction and purification

Lipopolysaccharides (LPS) were extracted using Apicella’s modification of the hot phenol method (48,49) as previously described (34). LPS extracts were purified by adding 50% aqueous trichloroacetic acid (CCl_3_CO_2_H) at 4°C to precipitate proteins and nucleic acids. The supernatant was then dialyzed against distilled water and lyophilized to yield the corresponding LPSs. All LPS samples were checked for contaminating proteins, nucleic acids and other lipid classes as previously described (34). Each LPS fraction was assessed for metamorphic induction in larvae of *H. elegans* at 5 µg·ml^-1^ and 10 µg·ml^-1^.

### Separation of O-antigen and Lipid A components

O-antigens were separated from lipid A components through mild acid hydrolysis of the LPSs with 2% aqueous acetic acid at 100°C until lipid precipitation (6 h). The precipitate was pelleted by centrifugation (13,000 × g, 20 min), and the resulting supernatant was collected and lyophilized for use in settlement and metamorphosis assays. Spectrographic analysis confirmed that O-antigen samples were free of contaminating lipids and lipid A samples were free of contaminating polysaccharides (50).

### Settlement assays

Settlement assays with larvae of *P. acuta* were conducted in 12-well plates. Each replicate evaluated 10 larvae, with 6 replicates per assay. Assays were performed in three independent trials to ensure reproducibility. Larval competency was confirmed through the use of the artificial inducer 5 mM caesium chloride or wild biofilms as positive controls (20). Wild biofilms were generated by submerging calcium carbonate plugs in the field for over one month, allowing natural colonization and maturation of microbial and algal communities on the surface. Sterile seawater (DFASW) was used as the negative control. Settlement and metamorphosis were counted after 12 and 24 h exposure to biofilms. Because settlement and metamorphosis can occur independently in coral larvae, metamorphosis was recorded separately from settlement (22). Larvae that metamorphosed at the air–water interface were classified as metamorphosed but unattached, while those that attached to the experimental surface and then metamorphosed were recorded as metamorphosed and attached.

### Commercial monosaccharides

The monosaccharides D-glucose, D-mannose, and D-glucuronic acid were acquired from Sigma Aldrich. Individual stock solutions of each monosaccharide were made (10 mM) in sterile seawater and diluted to experimental concentrations (1 mM and 0.1 mM) before addition of larvae. Larvae were then observed for settlement and metamorphosis as described above.

### Lectin Treatments

Monospecific biofilms of *T. euphylliae* M23b were prepared as above. Prior to addition of larvae for settlement assays, replicate biofilms were treated with the lectins Concanavalin A (ConA) and Wheatgerm Agglutinin (WGA). ConA is mannose binding and WGA is N-acetylglucosamine and sialic acid binding. Both mannose and N-acetylglucosamine have been implicated in other marine invertebrate metamorphosis studies (13,51,52). Additional controls of sterile seawater with each lectin were included.

### Statistical analysis

All statistical analyses were performed in Graphpad Prism 9 for Windows, (GraphPad Software, San Diego, California USA, www.graphpad.com). Significant differences (p<0.05) were calculated for total metamorphosis using Kruskal-Wallis followed by Dunn’s test for multiple comparisons with false detection rate (FDR) correction (53).

### Data availability

All data used in this manuscript are deposited in Figshare and are publicly available https://figshare.com/s/97d4324228ce4842ee1d

## Results

### Monospecific biofilms

Larvae of *P. acuta* were induced to settle and metamorphose by monospecific biofilms from *P. luteoviolacea* B1P, but not from the strains HI1 and ATCC 33492. Similarly, larvae of *P. acuta* were induced to settle and metamorphose by monospecific biofilms from *T. euphylliae* M23b, but not from *T. euphylliae* MR31e. Neither *C. lytica* HI1 nor *T. aiptasiae* T48 induced settlement and metamorphosis in the coral larvae (Figure 1A). Metamorphosis without attachment was only observed for P. luteoviolacea B1P biofilms (Figure 1A).

**Figure 1.**
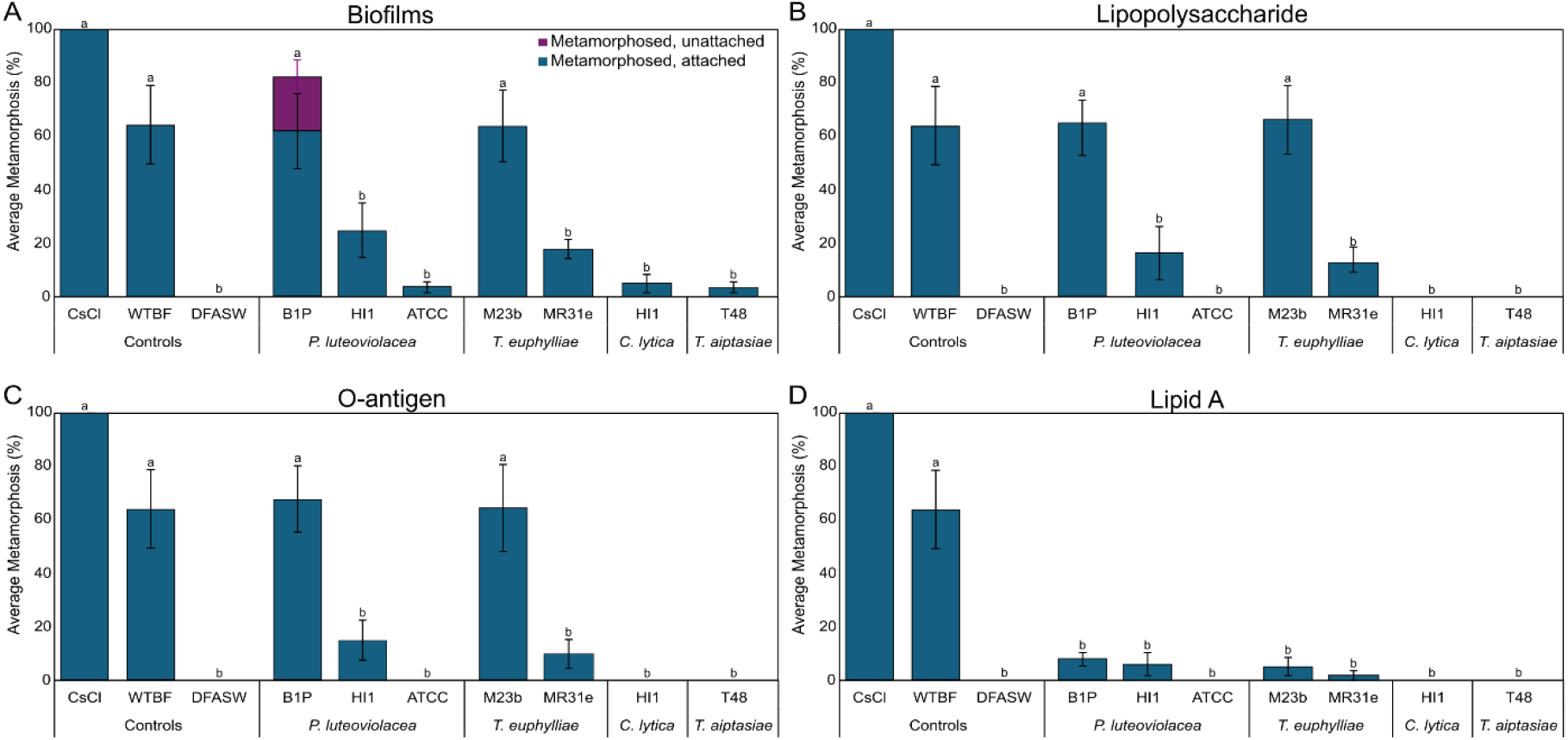
Settlement and metamorphosis in larvae of *Pocillopora acuta* after 24 h exposure to A) monospecific bacterial films, B) purified LPS, C) O-antigen, and D) Lipid A. Wild type biofilms (WT BF) and caesium chloride (5mM CsCl) were used as positive controls and sterile seawater (DFASW) was used for negative controls. Plots are averaged from three independent settlement bioassays. Error bars represent standard error. Letters indicate significant differences in total metamorphosis calculated with Kruskal-Wallis and Dunn’s test.

### LPS extracts

Larvae of *P. acuta* were induced to settle and metamorphose by LPS extracted from *P. luteoviolacea* B1P, but not P. luteoviolacea HI1 or ATCC 33492. Larvae of *P. acuta* were induced to settle and metamorphose by LPS extracted from *T. euphylliae* M23b, but not *T. euphylliae* MR31e. LPS extracted from *C. lytica* HI1 or *T. aiptasiae* T48 was not inductive (Figure 1B).

### O-antigen and lipid A samples

*P. acuta* were induced to settle and metamorphose by isolated O-antigens from *P. luteoviolacea* B1P and *T. euphylliae* M23b only (Figure 1C). No lipid A samples induced metamorphosis in these larvae (Figure 1D).

### Impact of lectins on inductive activity of *T*. *euphylliae*

The metamorphic inductive activity of *T. euphylliae* M23b was not inhibited by pretreatment with Con A but was partially inhibited by treatment with WGA (Figure 2).

**Figure 2.**
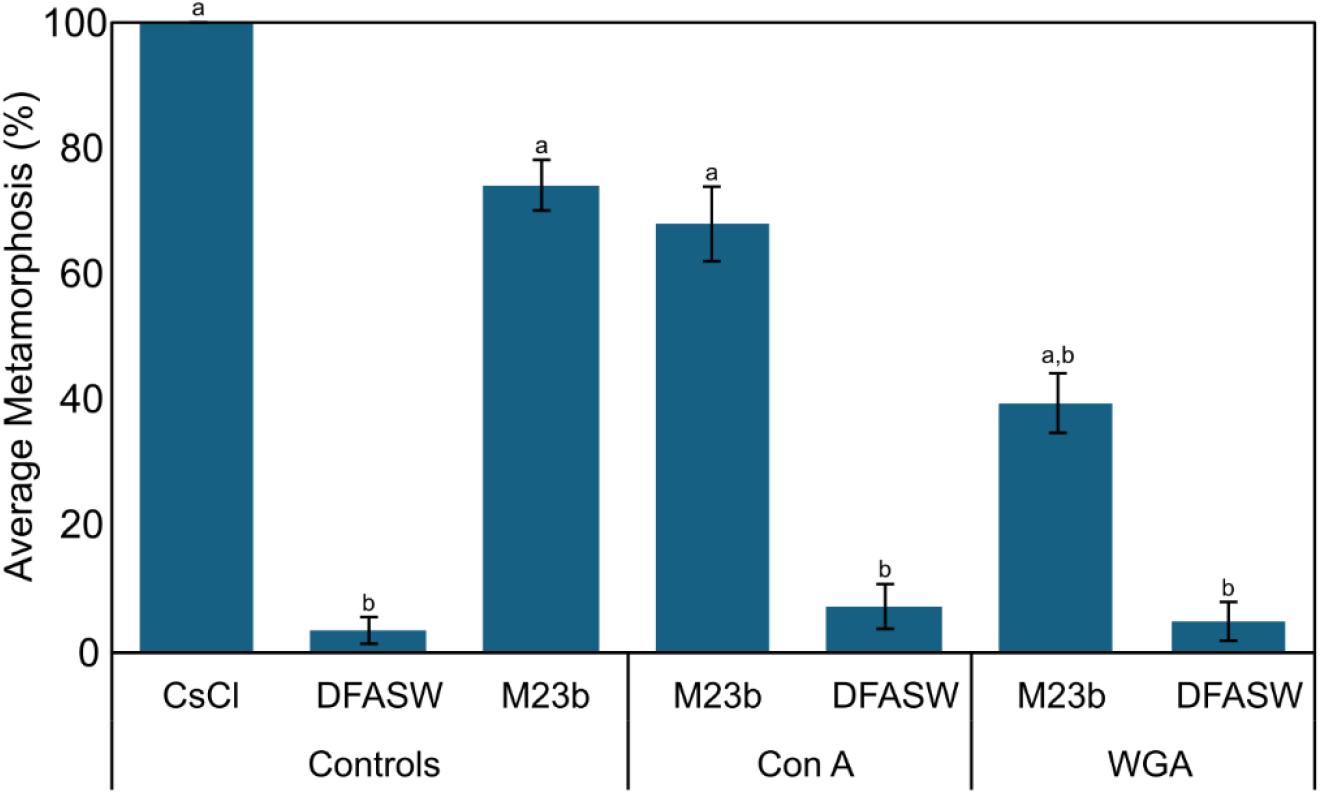
Settlement and metamorphosis in larvae of *P. acuta* after 24 h exposure to monospecific biofilms of *T. euphylliae* M23b with or without a 1 hr preexposure with the lectins Concanavalin A (ConA) or Wheatgerm agglutinin (WGA). Positive larval control: caesium chloride (5mM CsCl), Positive experimental control: biofilms of *T. euphylliae* M23b. Negative control: sterile seawater (DFASW). Plots are average from three independent settlement bioassays. Error bars represent standard error. Letters indicate significant differences in total metamorphosis calculated with Kruskal-Wallis and Dunn’s test.

### Impact of monosaccharides on inductive activity of *T*. *euphylliae*

Individual monosaccharides did not recreate the activity of the isolated O-antigen or monospecific biofilms of *T. euphylliae* M23b (Figure 3).

**Figure 3.**
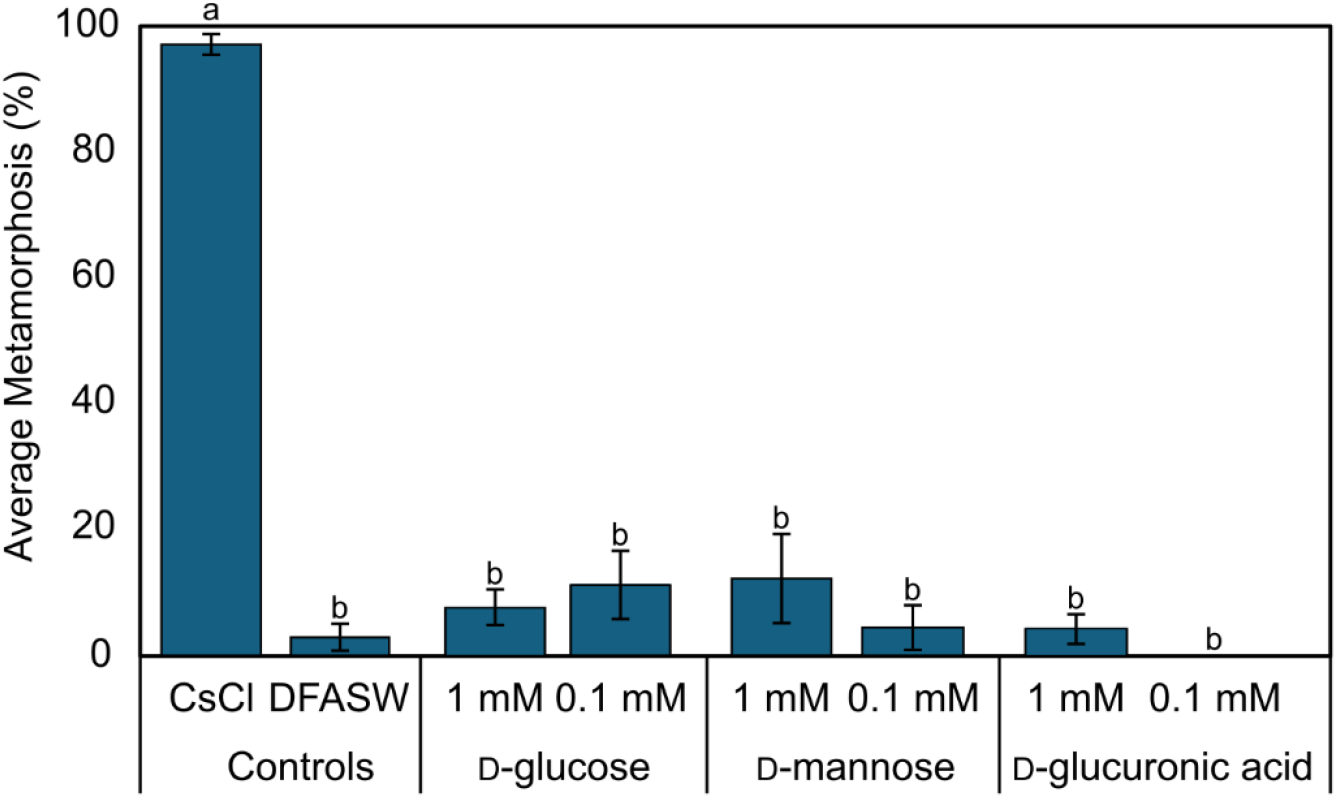
Settlement and metamorphosis in larvae of *Pocillopora acuta* after 24 h exposure to commercial monosaccharides (1mM or 0.1mM). Positive larval control: caesium chloride (5 mM CsCl). Negative control: sterile seawater (DFASW). Plots are average from three independent settlement bioassays. Error bars represent standard error. Letters indicate significant differences in total metamorphosis calculated with Kruskal-Wallis and Dunn’s test.

## Discussion

Here, we demonstrate that O-antigen from purified lipopolysaccharide (LPS) derived from specific, inductive bacterial strains can directly induce metamorphosis in larvae from the coral *Pocillopora acuta*. This result aligns with prior demonstrations of LPS-mediated settlement in the serpulid tubeworm *Hydroides elegans* (34). That the same class of molecules can induce these two organisms despite their deep evolutionary divergence suggests these molecules are either a conserved or repeatedly convergently evolved mechanism/cue regulating this pivotal life-history transition.

O-antigens are uniquely suited for a role in the induction of metamorphosis due to their immense structural complexity (54–58). Unlike nucleic acids and proteins, which are synthesized via template-based mechanisms, glycan biosynthesis proceeds through non-template-driven enzymatic pathways, yielding branching architectures, variable monosaccharide compositions, diverse linkages, and anomeric configurations (54,55). This resultant combinatorial potential, often referred to as the “glycan code,” enables highly specific biological messages to be encoded in glycan structure (54,55). In the context of larval settlement, this modularity allows glycans to serve as environmentally and evolutionarily adaptable signals where subtle structural differences can yield distinct biological outcomes. This specificity is evident in our experimental findings: only a subset of O-antigen preparations induced metamorphosis in *P. acuta*, and induction was strain-specific, even among bacteria of the same species. Furthermore, when inductive bacterial species were shared between the coral and tubeworm species, typically occupying different habitats, the inductive strains did not overlap.

That induction of metamorphosis by O-antigen differed not only between invertebrate larval species but also between bacterial strains of the same bacterial species suggests the potential ability of invertebrate larvae to discriminate among habitats based on fine-scale glycan structures, possibly via lectin-like receptors on their surfaces (37). This idea is supported by early experimental work on the polychaete *Janua (Dexiospira) brasiliensis*, where larval settlement and metamorphosis were shown to be inhibited by the monosaccharide D-glucose and by enzymatic treatments that disrupt glycoprotein structure, indicating a lectin-mediated recognition system between larval surface proteins and bacterial exopolysaccharides (37). In this hypothesis, lectins on the surface of the larva recognize and bind to glucose-containing sugar structures (glycoconjugates) found in the bacterial exopolymer. This binding triggers metamorphosis only when the shape and structure of the sugar molecules are compatible with the lectins, suggesting that a precise molecular match is required for the response to occur Such a recognition system would enable larvae to interpret microbial cues as proxies for habitat suitability during substrate selection. This highlights a critical point: the ability of larvae to distinguish subtle glycan variants may underlie species-specific habitat selection.

For a metamorphic cue to be ecologically relevant, it must be presented in a spatially constrained and biologically interpretable manner (59). O-antigens fulfil this criterion, because they are surface-bound, stable and localized to microbial biofilms, in contrast to diffusible, waterborne cues that provide ambiguous spatial information (59). This spatial localization ensures that larvae encounter cues only upon direct surface contact, allowing for fine-scale habitat discrimination and reducing the risk of inappropriate or premature metamorphosis (59,60). Our observations show that O-antigen induced metamorphosis in *P. acuta* is dose-dependent, consistent with the estimated concentrations of LPS in natural biofilms. These findings support the ecological relevance of O-antigen as a substrate-linked metamorphic cue. Future work examining structure–activity relationships, particularly through selective modification or removal of O-antigen moieties, will help define the specific glycan features responsible for inductive activity.

O-antigen is not the first or only surface-bound glycan to be identified as inductive for metamorphosis in marine invertebrate larvae. Indeed, surface-bound glycans are emerging as a unifying class of metamorphic inducers across diverse marine invertebrates. A growing body of evidence shows that structurally complex, surface-associated glycans can act as potent and ecologically relevant cues for larval settlement in taxa as diverse as annelids (34,61), molluscs (36,62,63), and cnidarians (13,16,52,64). For example, bacterial exopolysaccharides such as Rha-Man and curdlan induce settlement in the commensal hydrozoan *Hydractinia echinata* (52), while in *acroporid* corals, glycosylated lipids and polysaccharides from crustose coralline algae function as settlement signals (13,16,64). These chemically distinct but functionally similar molecules highlight a common biochemical motif—persistent, substrate-linked glycans—that facilitate species-specific recognition of suitable habitats. This specificity supports the notion of a modular, evolutionarily adaptable system that enables larvae to interpret complex landscapes during substrate selection.

In natural reef environments, coral larvae are unlikely to encounter monospecific bacterial biofilms or crustose coralline algae (CCA) devoid of a microbial coat. Instead, they interact with chemically complex, multispecies biofilms shaped by substrate type, hydrodynamics, and environmental gradients (10,24,65). Biofilms that develop on CCA or macroalgae differ markedly in microbial composition and metabolic output from those on bare rock or rubble (65–67). Even among CCA species, inductive and non-inductive biofilms support distinct microbial communities (68). This heterogeneity creates a chemically complex and temporally dynamic landscape that larvae must navigate. This ecological complexity underscores the need to consider settlement cues not in isolation but as part of a broader, multimodal sensory environment (12). This hypothesis aligns with broader patterns of cue synergy observed in marine invertebrate settlement and provides a mechanistic explanation for the variable effects of cues across taxa and conditions.

This study provides mechanistic insight into how larvae interpret complex microbial landscapes and points to the evolutionary recruitment of bacterial surface glycans as reliable indicators of habitat suitability. Understanding the molecular basis of glycan-mediated signalling opens the door to targeted reef restoration strategies. It should be noted, however, that the very molecular features that make surface-bound glycans ideal cue molecules, their variability and in the case of O-antigen, their sensitivity to environmental conditions also signals the vulnerability of these interactions to environmental disruptions. Anthropogenic stressors such as ocean warming, acidification, and pollution threaten the microbial communities that produce inductive cues and altered microbial community dynamics may shift the availability of inductive taxa and the molecules that they produce (24,69).

Despite identifying LPS as a potent cue, important questions remain. The larval receptor systems that detect LPS, and the precise moieties within O-antigens responsible for activity remain to be resolved. The observed variability in cue effectiveness among closely related bacterial strains reinforces the understanding that larval sensory systems are finely tuned to discriminate among glycans at high resolution. Future studies should investigate how o-antigens interact with other known cues—such as algal-derived glycans and small hydrophobic bacterial metabolites—and whether these cues act synergistically, additively, or hierarchically in driving settlement. Comparative analyses across coral species and *in situ* validations will be essential to establish the generality and ecological relevance of glycan-based settlement mechanisms.

